# Complete genome sequence of *Bacillus velezensis* JT3-1, a microbial germicide isolated from yak feces

**DOI:** 10.1101/555219

**Authors:** Youquan Li, Xuan Li, Dan Jia, Junlong Liu, Jinming Wang, Aihong Liu, Zhijie Liu, Guiquan Guan, Guangyuan Liu, Jianxun Luo, Hong Yin

## Abstract

*Bacillus velezensis* JT3-1 is a probiotic strain isolated from feces of the domestic yak (*Bos grunniens*) in the Gansu province of China. It has strong antagonistic activity against *Listeria monocytogenes*, *Staphylococcus aureus*, *Escherichia coli, Salmonella* Typhimurium, *Mannheimia haemolytica*, *Staphylococcus hominis*, *Clostridium perfringens*, and *Mycoplasma bovis*. These properties have made the JT3-1 strain the focus of commercial interest. In this study, we describe the complete genome sequence of JT3-1, with a genome size of 3,929,799 bp, 3761 encoded genes and an average GC content of 46.50%. Whole genome sequencing of *Bacillus velezensis* JT3-1 will lay a good foundation for elucidation of the mechanisms of its antimicrobial activity, and for its future application.

---

Antibiotics utilized in animal husbandry introduce various problems while improving animal productivity (Sharma et al., 2017). In many countries, the current tendency in animal production is towards a reduction or prohibition of the use of antibiotics, with an increase in application of non-antibiotic methods (Lillehoj and Lee, 2012). Of the non-antibiotic methods, dietary supplementation of can improve the productivity and immunological status of livestock (Abd El-Tawab et al., 2016; Khan et al., 2016; Reuter, 2001) Probiotics are dietary supplements that are by definition "live microorganisms, which, upon ingestion in sufficient numbers, exert health benefits" (Schrezenmeir and de Vrese, 2001). In ruminants, probiotics can change the rumen microbial ecosystem, and promote nutrient digestibility and feed efficiency. The most commonly used probiotics are *Lactobacillus*, *Bifidobacterium*, *Bacillus*, and yeast strains. Supplemental *B. subtilis* increases growth performance, disease resistance and immune response in animals (Guo et al., 2016, 2017). Previous reports have demonstrated that *Bacillus velezenis* QST713 (Pandin et al., 2018), *Bacillus velezenis* M75 (Kim et al., 2017), and *Bacillus velezenis* S3-1 (Jin et al., 2017) showed a strong ability to form biofilm and inhibit pathogens.

*B. velezensis* JT3-1 was isolated from the feces of domestic yak (*Bos grunniens*) in Gansu Province, China. Our study showed that the *B. velezensis* JT3-1 strain had strong antagonistic activity against various intestinal pathogenic flora, and stimulation of animal (such as sheep, goat, mouse, pig and cattle) growth (not published). However, there are no published reports on the novel *B. velezensis* JT3-1 strain and its genome sequence is not found in Genebank. In this study, the genome sequence of the novel strain JT3-1 was determined, the secondary metabolite clusters in the genome of *B. velezensis* JT3-1 were analyzed, and an antimicrobial activity assay was performed to investigate the abilities of the biocontrol strain.

## 1. General genomic features of *Bacillus velezensis* JT3-1

Genomic DNA was extracted using Gentra Puregene Yeast/Bact (Qiagen, Germany), and the whole-genome sequencing of *B. velezensis* JT3-1 was performed using a PacBio Sequel sequencing platform at Beijing Genewiz Bioinformatics Technology Co., Ltd. A library with a 10 kb insert size was constructed for the PacBio Sequel platform, and the library was sequenced in a PacBio SMRT (Chin et al., 2013) instrument. Subsequently, the PacBio reads were assembled using SmrtLink, and a total of 199380 reads, in about 550 Mb, were obtained. Finally, we obtained a single circular gapless chromosome. Prodigal software (Hyatt et al., 2010) was utilized to find coding genes in the bacterium. Transfer RNAs (tRNAs) were detected in the genome using the program blast based on Rfam (Eric et al., 2014). The coding genes were annotated with the National Center for Biotechnology Information (NCBI) nr database by Diamond (Buchfink et al., 2015). Afterwards, the functions of genes were annotated using the Gene Ontology database, and the pathways were annotated using the Kyoto Encyclopedia of Genes and Genomes database. The proteins encoded by the genes were phylogenetically classified by the Clusters of Orthologous Groups database. The potential genes involved with the biosynthesis of bacteriocins were identified with antiSMASH4.0 (Blin et al., 2017) (http://antismash.secondarymetabolites.org/). The complete genome of *B. velezensis* JT3-1 contains one gapless circular chromosome of 3,929,799 bp, 3761 protein encoding genes, 86 tRNA genes, 90 rRNA genes and 83 other ncRNA; it shows a G + C content of 46.50%, and no plasmids were found (**Table 1**). The circular genome map of the *B. velezensis* JT3-1 complete genome is displayed in **Fig. 1**.

**Table 1.** General genome features of *Bacillus velezensis* JT3-1.

| Feature | Value |
| --- | --- |
| Genome size (bp) | 3,929,799 |
| GC content [%] | 46.50 |
| Predicted genes | 3761 |
| Total ncRNA | 259 |
| tRNA genes | 86 |
| rRNA genes | 90 |
| Other ncRNA | 83 |

## 2. Antimicrobial activity of *Bacillus velezensis* JT3-1

The antimicrobial activity of *Bacillus velezensis* JT3-1 was investigated; an agar well diffusion assay was utilized to test the antibacterial properties of the JT3-1 culture supernatant, as previously described (Liu et al., 2017). The Nutrient Agar (NA) medium was utilized to culture the remaining bacteria, and the PPLOB medium was utilized to culture *Mycoplasma bovis*. The diameter of the inhibition zones was calculated in millimeters, and the plus sign shows the range of the inhibition zone. The three-plus sign (+++) means that the diameter of the inhibition zone is more than 15 mm; the two-plus sign (++) means that the diameter of the inhibition zone is between 10 mm and 15 mm; and the one-plus sign (+) means that the diameter of the inhibition zone is less than 10 mm (the diameter of the wells was 5 mm). Each test was repeated three times, and the mean values were adopted. Our study identified that *B. velezensis* JT3-1 showed strong antimicrobial activity against some Gram-negative human or animal pathogens (such as *E. coli* ATCC43888, *E. coli* ATCC25922, *E. col*i LQ1-2, *Yersinia enterocolitis* 23715, *Proteus vulgaris* ATCC29905, *Shigella bogdii* ATCC9207, *Salmonella* Typhimurium ATCC13311, and *Mannheimia haemolytica* ATCC29696), several Gram-positive animal and human pathogenic bacteria (such as *Streptococcus pyogenes* ATCC19615, *Listeria monocytogenes* ATCC19115, *Staphylococcus hominis* ZSY2, *Staphylococcus aureus* ATCC6538, and *Clostridium perfringens* MQF5), and *Mycoplasma bovis* CYF strain (**Table 2**). Our previous study identified that the *B. velezensis* JT3-1 strain plays an important role in preventing and curing calf and lamb diarrhea caused by bacteria or by the weaning period of young animals (not published).

**Table 2.** Secondary metabolite clusters identified in the genome of *B. velezensis* JT3-1.

| Pathogens | Broth medium | Antimicrobial activity |
| --- | --- | --- |
| Gram-negative bacteria |  |  |
| *E.coli* ATCC43888 | NA<sup>a</sup> | +++ |
| *E.coli* ATCC25922 | NA | +++ |
| *E.coli* LQ1-2 | NA | +++ |
| *Yersinia enterocolitis* 23715 | NA | ++ |
| *Proteus vulgaris* ATCC29905 | NA | + |
| *Shigella bogdii* ATCC9207 | NA | + |
| *Salmonella typhimurium* ATCC13311 | NA | ++ |
| *Mannheimia haemolytica* ATCC29696 | NA | ++ |
| Gram-positive bacteria |  |  |
| *Listeria monocytogenes* ATCC19115 | NA | +++ |
| *Staphylococcus aureus* ATCC6538 | NA | +++ |
| *Streptococcus pyogenes* ATCC19615 | NA | +++ |
| *Staphylococcus hominis* ZSY2 | NA | +++ |
| *Clostridium perfringens* MQF5 | NA | +++ |
| *Mycoplasma bovis* CYF | PPLOB<sup>b</sup> | +++ |

<sup>a</sup> NA: Nutrient Agar.<sup>b</sup> PPLOB: PPLOB medium was utilized to culture *Mycoplasma bovis*.

## 3. Secondary metabolite clusters

Through the antiSMASH 4.0 genome analysis tool (Blin et al., 2017), 30 clusters of secondary metabolites were identified in the genome of the JT3-1 strain: two transATPKS-NRPS, two transATPKS (trans-acyl transferase polyketide synthetase), one lantipeptide, one encoding NRPS (non-ribosomal peptide synthetase), one T3PKS, one bacteriocin-Nrps, two terpenes, one other KS, three Cf_fatty_acid, five Cf_saccharide, and 11 Cf_putative (**Table 3**). Eight clusters were clearly involved in the synthesis of difficidin, surfactin, fengycin, butirosin, macrolactin, bacilysin, bacillaene and bacillibactin, and they were closely related with antimicrobial activities (Chen et al., 2009; Niazi et al., 2014; Ongena et al., 2005; Liu et al., 2017; Kim et al., 2017; Stein, 2005; Pandin et al., 2018). This analysis showed that at least 17% of the JT3-1 genome contributed to the transport and regulation of antibiotic biosynthesis.

**Table 3.**
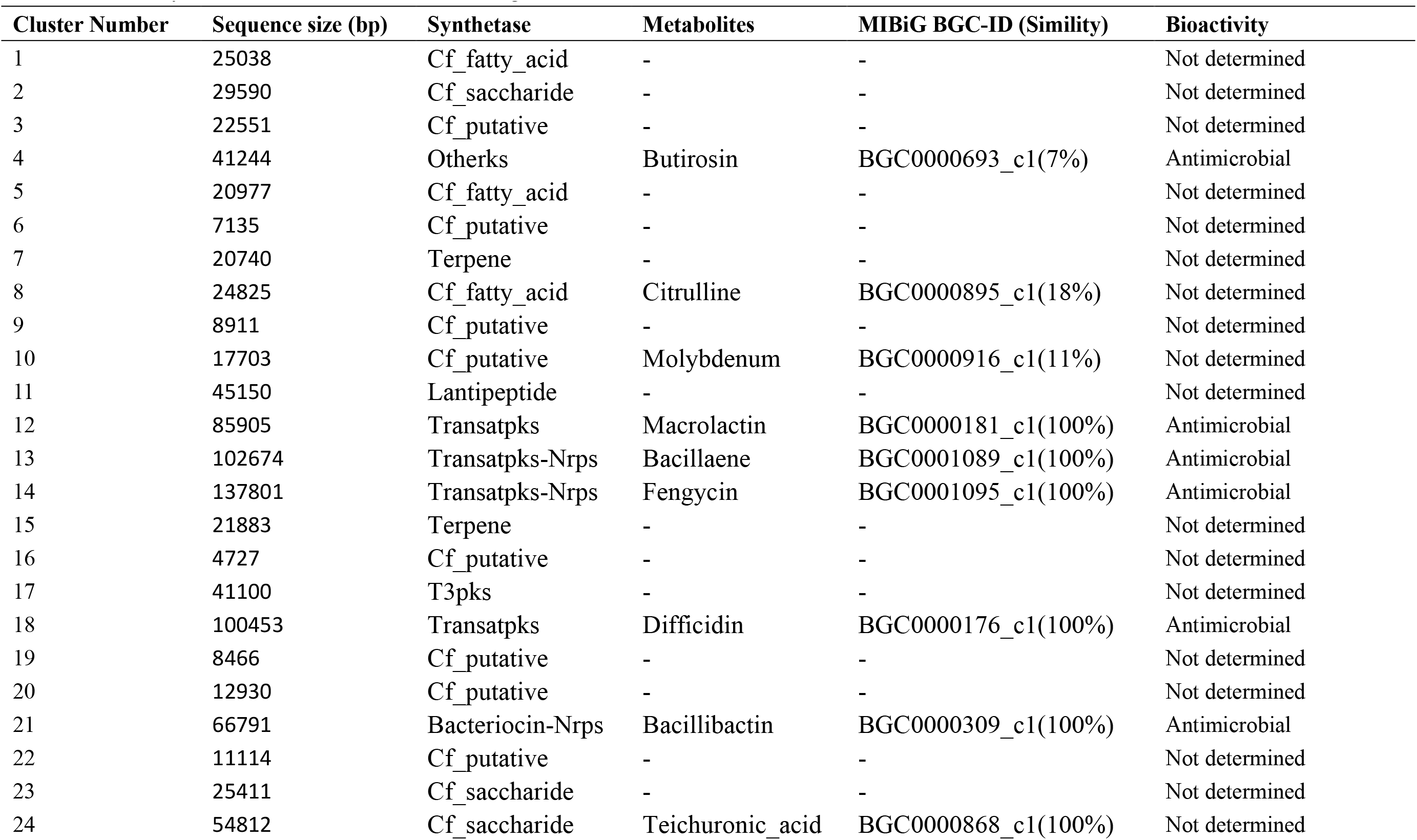
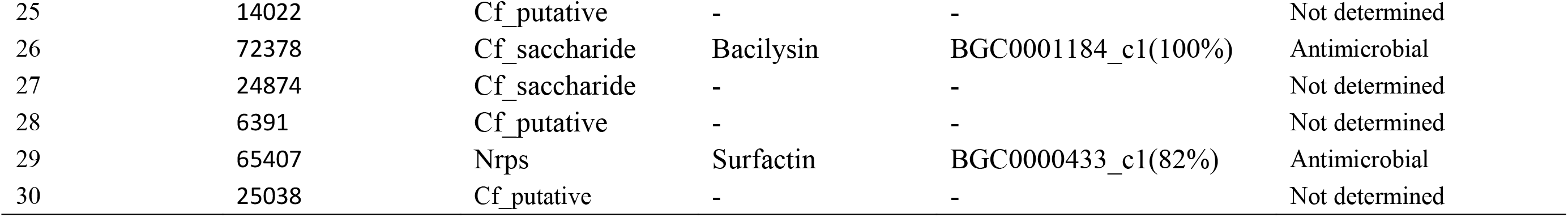
Antimicrobial activity of the *Bacillus velezensis* JT3-1 strain. ^a^ NA: Nutrient Agar. ^b^ PPLOB: PPLOB medium was utilized to culture *Mycoplasma bovis*.

| Cluster Number | Sequence size (bp) | Synthetase | Metabolites | MIBiG BGC-ID (Simility) | Bioactivity |
| --- | --- | --- | --- | --- | --- |
| 1 | 25038 | Cf_fatty_acid | - | - | Not determined |
| 2 | 29590 | Cf_saccharide | - | - | Not determined |
| 3 | 22551 | Cf_putative | - | - | Not determined |
| 4 | 41244 | Otherks | Butirosin | BGC0000693_c1(7%) | Antimicrobial |
| 5 | 20977 | Cf_fatty_acid | - | - | Not determined |
| 6 | 7135 | Cf_putative | - | - | Not determined |
| 7 | 20740 | Terpene | - | - | Not determined |
| 8 | 24825 | Cf_fatty_acid | Citrulline | BGC0000895_c1(18%) | Not determined |
| 9 | 8911 | Cf_putative | - | - | Not determined |
| 10 | 17703 | Cf_putative | Molybdenum | BGC0000916_c1(11%) | Not determined |
| 11 | 45150 | Lantipeptide | - | - | Not determined |
| 12 | 85905 | Transatpks | Macrolactin | BGC0000181_c1(100%) | Antimicrobial |
| 13 | 102674 | Transatpks-Nrps | Bacillaene | BGC0001089_c1(100%) | Antimicrobial |
| 14 | 137801 | Transatpks-Nrps | Fengycin | BGC0001095_c1(100%) | Antimicrobial |
| 15 | 21883 | Terpene | - | - | Not determined |
| 16 | 4727 | Cf_putative | - | - | Not determined |
| 17 | 41100 | T3pks | - | - | Not determined |
| 18 | 100453 | Transatpks | Difficidin | BGC0000176_c1(100%) | Antimicrobial |
| 19 | 8466 | Cf_putative | - | - | Not determined |
| 20 | 12930 | Cf_putative | - | - | Not determined |
| 21 | 66791 | Bacteriocin-Nrps | Bacillibactin | BGC0000309_c1(100%) | Antimicrobial |
| 22 | 11114 | Cf_putative | - | - | Not determined |
| 23 | 25411 | Cf_saccharide | - | - | Not determined |
| 24 | 54812 | Cf_saccharide | Teichuronic_acid | BGC0000868_c1(100%) | Not determined |
| 25 | 14022 | Cf_putative | - | - | Not determined |
| 26 | 72378 | Cf_saccharide | Bacilysin | BGC0001184_c1(100%) | Antimicrobial |
| 27 | 24874 | Cf_saccharide | - | - | Not determined |
| 28 | 6391 | Cf_putative | - | - | Not determined |
| 29 | 65407 | Nrps | Surfactin | BGC0000433_c1(82%) | Antimicrobial |
| 30 | 25038 | Cf_putative | - | - | Not determined |

## 4. Genes involved in promoting animal growth

Besides producing many secondary metabolites with antimicrobial activity, several *Bacillus* spp. possess high secretory ability and some strains utilize ''cell factories" to produce industrial enzymes (Ruiz-Garcia et al., 2005; Niazi et al., 2014). Several *B. subtilis* strains can colonize the intestinal microvilli of the animal, support nutrition, and trigger a healthy immune system (Guo et al., 2016, 2017; Abd El-Tawab et al., 2016). Several strains of *B. amyloliquefaciens* subsp. *plantarum* can improve plant growth and confer protection by producing antimicrobial materials and phytohormones (Chen et al., 2006; Koumoutsi et al., 2004). *B. velezensis* strains are able to suppress plant or animal pathogens, increase plant or animal growth, and very efficiently colonize rhizosphere soil and the intestinal tract of the host (Ruiz-Garcia et al., 2005; Jin et al., 2017; Chen et al., 2017; Wang et al., 2018; Yi et al., 2018). Our previous study showed that *B. velezensis* JT3-1 can efficiently colonize the intestinal microvilli of an animal, promote growth, and trigger the immune system (not published). The genome of the *B. velezensis* JT3-1 strain contains many genes involved in biofilm formation and animal intestinal colonization, including the *als*D, *bdh*A, *sac*B, *xyn*A, *xyn*D, 3-hydroxy-2-butanone, 2,3-butanediol, Bacllibactin, indole acetic acid, trehalose, and phytase genes. Several genes of *B. velezensis* JT3-1 were related to the enzymes encoded to use derived substrates; *xyn*D and *xyn*A, which encode xylanase, are related to the use of hemicelluloses and celluloses presented in the animal intestinal tract. *B. velezensis* JT3-1 also harbors several genes involved in the synthesis of acetoin, 3-hydroxy-2-butanone and 2,3-butanediol (*bdh*A and *als*D), which are able to improve animal and plant growth efficiency and trigger systemic defenses against pathogenic agents. Bacillibactin, which is a nonribosomal peptide previously discovered in *B. subtilis*, can transfer iron to iron acquisition mechanisms that are essential to animal and plant life (Zhou et al., 2018). Trehalose is a non-reducing disaccharide that plays an important role in protecting cells from stress and maintaining cell structure (Wang et al., 2019). Phytase is secreted by the intestinal bacteria of ruminant animals (such as camels, deer, cattle, goats and sheep), and it can degrade phytate and improve the efficacy of phosphorus metabolism and the performance of animals and plants (Frias et al., 2003). At present, phytase is widely used as an animal feed supplement. Meanwhile, *B. velezensis* JT3-1 also harbors the genes that encode ATP-dependent Clp protease (*clp*E), α-glucosidase (*gly*A), β-glucosidase (*gmu*D), catalase (*kat*E), and thiol peroxidase (*tpx*), among others, demonstrating that *B. velezensis* JT3-1 is able to use many organic materials as nutritional supplies to promote its survivability.

Until now, 167 *B. velezensis* strains have been sequenced and released in the NCBI genome database. The availability of the complete genome sequence of *B. velezensis* JT3-1 will not only enrich the genome database but also provide some valuable information regarding the molecular mechanism of antimicrobial action and build the foundation for the utilization of JT3-1 as a biocontrol agent in animal husbandry.

## 5. Nucleotide sequence accession number

The complete genome sequence of *B. velezensis* JT3-1 was submitted to GeneBank under the accession number CP032506. This strain was deposited at the China General Microbiological Culture Collection Center under the accession number CGMCC No.15545.

## Acknowledgments

This study was supported by grants from the National Key R&D Program of China (2018YFD0501804, 2018YFD0502305 and 2017YFD0501200), MOST; NBCITS (CARS-37).

---

Fig. 1. Circular genome maps of the *Bacillus velezensis* JT3-1 complete genome. From the outermost circle to the inner, circle 1, the size of complete genome; circles 2 (red) and 3 (green), the predicted protein-encoding genes on the + and - strands, respectively; circle 4, rRNA/tRNA; circle 5, G + C ratio; circle 6, repetitive sequence.

## References

Abd El-Tawab, M.M., Youssef, I.M., Bakr, H.A., Fthenakis, G.C., Giadinis, N.D., 2016. Role of probiotics in nutrition and health of small ruminants. Pol. J. Vet. Sci. 19(4),893–906.

Blin, K., Wolf, T., Chevrette, M.G., Lu, X., Schwalen, C.J., Kautsar, S.A., Suarez, Duran, H.G., de Los Santos, E.L.C., Kim, H.U., Nave, M., Dickschat, J.S., Mitchell, D.A., Shelest, E., Breitling, R., Takano, E., Lee, S.Y., Weber, T., Medema, M.H., 2017. antiSMASH 4.0-improvements in chemistry prediction and gene cluster boundary identification. Nucleic Acids Res. 45(W1),W36–W41. 10.1093/nar/gkx319

Buchfink, B., Xie, C., Huson, D. H., 2015. Fast and sensitive protein alignment using DIAMOND. Nat. methods. 12(1), 59–60.

Chen, X.H., Vater, J., Piel, J., Franke, P., Scholz, R., Schneider, K., Koumoutsi, A., Hitzeroth, G., Grammel, N., Strittmatter, A.W., Gottschalk, G., Süssmuth, R.D., Borriss, R., 2006. Structural and functional characterization of three polyketide synthase gene clusters in Bacillus amyloliquefaciens FZB42. J Bacteriol 188: 4024–4036.

Chen, X.H., Koumoutsi, A., Scholz, R., Schneider, K., Vater, J., Süssmuth, R., Piel, J., Borriss, R., 2009. Genome analysis of Bacillus amyloliquefaciens FZB42 reveals its potential for biocontrol of plant pathogens. J. Biotechnol. 140, 27–37.

Chen, L., 2017. Complete genome sequence of Bacillus velezensis LM2303, a biocontrol strain isolated from the dung of wild yak inhabited Qinghai-Tibet plateau. J. Biotechnol. 251,124–127.

Chin, C.S., Alexander, D. H., Marks, P., Klammer, A. A., Drake, J., Heiner, C., Clum, A., Copeland, A., Huddleston, J., Eichler, E. E., Turner, S. W., Korlach, J., 2013. Nonhybrid, finished microbial genome assemblies from long-read SMRT sequencing data. Nat. methods. 10(6), 563–569.

Eric, P. N., Sarah, W. B., Alex, B., Jennifer D., Ruth Y. E., Sean R. E., Evan W. F., Paul P. G., Thomas A. J., John, T. and Robert, D. F., 2014. Rfam 12.0: updates to the RNA families database. Nucleic Acids Res. 10.

Frias, J., Doblado, R., Antezana, J. R., Vidal-Valverde, C. N., 2003. Inositol phosphate degradation by the action of phytase enzyme in legume seeds. Food Chemistry. 81 (2): 233.

Guo, M., Hao, G., Wang, B., Li, N., Li, R., Wei, L., Chai, T., 2016. Dietary Administration of Bacillus subtilis Enhances Growth Performance, Immune Response and Disease Resistance in Cherry Valley Ducks. Front. Microbiol. 7,1975.

Guo, M., Wu. F., Hao, G., Qi, Q., Li, R., Li, N., Wei, L., Chai, T., 2017. Bacillus subtilis Improves Immunity and Disease Resistance in Rabbits. Front. Immunol. 8,354.

Liu, G., Kong, Y., Fan, Y., Geng, C., Sun, M., 2017. Whole-genome sequencing of Bacillus velezensis LS69, a strain with a broad inhibitory spectrum against pathogenic bacteria. J. Biotechnol. 249,20–24

Jin, Q., Jiang, Q., Zhao, L., Su, C., Li, S., Si, F., Li, S., Zhou, C., Mu, Y., Xiao, M., 2017. Complete genome sequence of Bacillus velezensis S3-1, a potential biological pesticide with plant pathogen inhibiting and plant promoting capabilities. J. Biotechnol. 259,199–203.

Khan, R.U., Shabana, N., Kuldeep, D., Karthik, K., Ruchi, T., Mutassim, M.A., Alhidary, I.A., Zahoor, A., 2016. Direct-fed microbial: beneficial applications, modes of action and prospects as a safe tool for enhancing ruminant production and safeguarding health. Int. J. Pharm. 12,220–231.

Kim, S.Y., Lee, S.Y., Weon, H.Y., Sang, M.K., Song, J., 2017. Complete genome sequence of Bacillus velezensis M75, a biocontrol agent against fungal plant pathogens, isolated from cotton waste. J. Biotechnol. 241,112–115.

Koumoutsi A, Chen, X.H., Henne, A., Liesegang, H., Hitzeroth, G., Franke, P., Vater, J., Borriss, R., 2004. Structural and functional charaterization of gene clusters directing nonribosomal synthesis of bioactive cylic lipopeptides in Bacillus amyloliquefaciens FZB42. J. Bacteriol. 186: 1084–1096.

Hyatt, D., Chen, G. L., LoCascio, P. F., Larimer, F. W., Hauser, L. J., 2010. Prodigal: prokaryotic gene recognition and translation initiation site identification. BMC bioinformatics. 11.

Lillehoj, H.S., and Lee, K.W., 2012. Immune modulation of innate immunity as alternatives-to-antibiotics strategies to mitigate the use of drugs in poultry production. Poult. Sci. 91,1286–1291.

Muhammed, A.A., He, J., 2018. Use of probiotics and botanical extracts to improve ruminant production in the tropics: A review. Anim. Nutr. 4(3),241–249.

Niazi, A., Manzoor, S., Asari, S., Bejai, S., Meijer, J., Bongcam-Rudloff, E., 2014. Genome analysis of Bacillus amyloliquefaciens subsp. plantarum UCMB5113:) a rhizobacterium that improves plant growth and stress management. PLoS One 9, e104651.

Ongena, M., Jacques, P., Touré, Y., Destain, J., Jabrane, A., Thonart, P., 2005. Involvement of fengycin-type lipopeptides in the multifaceted biocontrol potential of Bacillus subtilis. Appl. Microbiol. Biotechnol. 69,29–38.

Pandin, C., Le, Coq, D., Deschamps, J., Védie, R., Rousseau, T., Aymerich, S., Briandet, R., 2018. Complete genome sequence of Bacillus velezensis QST713: A biocontrol agent that protects Agaricus bisporus crops against the green mould disease. J. Biotechnol. 278,10–19.

Reuter, G., 2001. Probiotics–possibilities and limitations of their application in food, animal feed, and in pharmaceutical preparations for men and animals. Berl. Münch. Tierärztl. Wochenschr. 114, 410–419.

Ruiz-Garcia, C., Bejar, V., Martinez-Checa, F., Llamas, I., Quesada, E., 2005. Bacillus velezensis sp. nov., a surfactant-producing bacterium isolated from the river Vélez in Málaga, southern Spain. Int. J. Syst. Evol. Microbiol. 55, 191–195.

Schrezenmeir, J., de Vrese, M., 2001. Probiotics, prebiotics, and synbiotics—approaching a definition. Am. J. Clin. Nutr. 73(2 Suppl),361S–364S.

Sharma, C., Rokana, N., Chandra, M., Singh, B.Pal., Gulhane, R.D., Gill, J.P.S., Ray, P., Puniya, A.K., Panwar, H., 2017. Antimicrobial Resistance: Its Surveillance, Impact, and Alternative Management Strategies in Dairy Animals. Front Vet. Sci. 4,237.

Stein, T., 2005. Bacillus subtilis antibiotics: structures, syntheses and specific functions. Mol. Microbiol. 56, 845–857.

Wang, B., Liu, G., Balamurugan, V., Sui, Y., Wang, G., Song, Y., Chang, Q., 2019. Apatite nanoparticles mediate intracellular delivery of trehalose and increase survival of cryopreserved cells. Cryobiolo. 86,103–110.

Wang, N., Li, P., Wang, M., Chen, S., Huang, S., Long, M., Yang, S., He, J., 2018. The Protective Role of Bacillus velezensis A2 on the Biochemical and Hepatic Toxicity of Zearalenone in Mice. Toxins (Basel). 10(11). pii: E449.

Zhou, M., Liu, F., Yang, X., Jin, J., Dong, X., Zeng, K.W., Liu, D., Zhang, Y., Ma, M., Yang, D., 2018. Bacillibactin and Bacillomycin Analogues with Cytotoxicities against Human Cancer Cell Lines from Marine Bacillus sp. PKU-MA00093 and PKU-MA00092. Mar. Drugs. 10,16(1).

Yi, Y., Zhang, Z., Zhao, F., Liu, H., Yu, L., Zha, J., Wang, G., 2018. Probiotic potential of Bacillus velezensis JW: Antimicrobial activity against fish pathogenic bacteria and immune enhancement effects on Carassius auratus. Fish Shellfish Immunol. 78,322–330.

